# Beta_2_-adrenoceptor agonist salbutamol increases leg glucose uptake and metabolic rate but not muscle glycogen resynthesis in recovery from resistance exercise of the quadriceps in lean young men

**DOI:** 10.1101/2021.04.30.442161

**Authors:** Johan Onslev, Martin Thomasson, Jørgen Wojtaszewski, Jens Bangsbo, Morten Hostrup

## Abstract

**Content:** Beta_2_-agonists evoke potent acute increases in peripheral glucose uptake and energy expenditure at rest. Exercise has been shown to blunt these effects. Whether this attenuation is extended into recovery from exercise is unknown.

**Objective:** To examine the effect of beta_2_-agonists on leg glucose uptake and leg metabolic rate in recovery from exercise.

**Design:** In a randomized, placebo-controlled, cross-over study using arteriovenous balance technique and analysis of thigh muscle biopsies we investigated the effect of 24mg oral salbutamol (a selective beta_2_-agonist) on leg glucose, oxygen, and lactate at rest, during exercise, and in recovery, as well as on muscle glycogen resynthesis.

**Participants:** Healthy, lean, young men (n=12).

**Results:** Leg glucose uptake tended to be two-fold higher at rest (0.22±0.12mmol/min, P=0.06). Accumulated leg glucose uptake was higher in recovery (21.1±6mmol, P=0.018) with salbutamol, but not during exercise. Leg oxygen uptake was 80% greater at rest (11±2.1mmol/min, P<0.01). Accumulated leg oxygen uptake was higher in recovery (1755±348mL, P<0.01) with salbutamol, but not during exercise. Muscle glycogen was lower with salbutamol 0.5h (109±25mmol/mg dry-weight, P<0.01) and 5h (101±19mmol/mg dry-weight, P<0.01) into recovery, suggestive of augmented glycogen utilization during exercise. There was no difference in glycogen resynthesis or glycogen synthase activity in the 5-hour recovery period with salbutamol.

**Conclusions:** These findings suggest that while resistance exercise confounds the augmentation of leg glucose uptake and metabolic rate induced by beta_2_-agonist at rest, this suppression is not conserved into recovery from exercise.

## Introduction

Beta_2_-adrenoceptor agonists (beta_2_-agonists) are widely used as first-line treatment in chronic obstructive pulmonary disease and asthma [1,2]. While their main therapeutic application is to induce bronchodilation, beta_2_-agonists induce numerous effects due to the abundance of beta_2_-adrenoceptors in several tissues [3,4]. The role of beta_2_-adrenoceptors in skeletal muscle has attracted interest in pharmaceutical drug development because of their involvement in the regulation of myocellular metabolism and glucose uptake [5–7]. Indeed, beta_2_-agonists can have profound effects on muscle glucose metabolism [8,9], which is substantiated by the observation that femoral infusion of selective agonist terbutaline almost doubled resting leg glucose uptake and clearance of lean young men [8,9]. Furthermore, beta_2_-agonists increase energy expenditure – partly due to an increased metabolic rate of skeletal muscle, which, together with observations from cell lines and rodents showing promising effects of beta_2_-agonist on glucose uptake and tolerance [10,11], underline a therapeutic potential of beta_2_-agonists in metabolic diseases., such as type 2 diabetes (T2DM) and obesity [10,12].

A notable phenomenon, however, is that the effect of beta_2_-agonist on metabolic rate and peripheral glucose uptake is blunted by exercise [8,9,13,14]. In fact, the greater resting leg glucose clearance induced by beta_2_-agonist was even shown to shift to a lowered clearance during exercise compared to placebo [9]. Given that exercise is emphasized by official guidelines as first-line treatment in T2DM and obesity [15,16], it is relevant to assess whether the confounding effect of exercise on the beta_2_-adrenergic-mediated augmentation of metabolic rate and peripheral glucose uptake is extended into the recovery from exercise. If this is the case, the application of beta_2_-agonist as a means to lose weight and augment glucose disposal in metabolic diseases would be less favorable.

Beta_2_-agonists have been shown to increase glycogen utilization during exercise [14]. Glycogen depletion is associated with a more prominent activation of AMP-activated protein kinase (AMPK) [17], which is fundamental in the regulation of myocellular glucose uptake in recovery from exercise [18]. Accordingly, it is conceivable that beta_2_-agonists, while not known to impact muscle AMPK directly, may lead to greater activation of AMPK in this setting, hence facilitating glucose disposal in recovery of the exercised muscles. Furthermore, a potentially larger muscle glucose uptake induced by beta_2_-agonist will likely cause either increased incorporation of glucose into glycogen or a greater glycolytic flux. Given that such an increase in flux may exceed the rate for which pyruvate is transformed to Acetyl-CoA by pyruvate dehydrogenase (PDH), this will possibly lead to higher lactate production. Thus, it is relevant to examine whether beta_2_-agonist modifies lactate release of the exercised leg and PDH activation in recovery from exercise. Indeed, phosphorylation of PDH has been shown to be higher at rest with beta_2_-agonist [14] suggestive of a decreased PDH activity. Glycogen synthase (GS) is the rate-limiting enzyme in glycogenesis making its *in vitro* activity a good indicator of glycogen resynthesis in recovery from exercise. While an increased glycogenolysis during exercise would favor increased GS-activity due to lower glycogen levels [17], unselective adrenergic stimulation is known to inhibit the augmented activity of GS facilitated by insulin [19]. Thus, prediction of the effect of beta2-adrenergic stimulation in recovery from exercise on GS-activity and glycogen resynthesis is difficult and likely dependent on a balance between multiple cellular pertubations.

Herein, we investigated whether a potential confounding effect of resistance exercise on the beta_2_-adrenergic induced augmentation of metabolic rate and peripheral glucose uptake would diminish in recovery from exercise. Furthermore, we examined how selective beta_2_-agonism affected glycogen resynthesis and canonical signaling pathways in recovery of the exercised muscle. We hypothesized that the acute effects of beta_2_-agonists on leg glucose uptake and leg oxygen uptake observed at rest would be blunted during exercise, but restored in recovery accompanied by less muscle glycogen resynthesis.

## Methods

### Human subjects

Thirteen healthy trained young men volunteered to participate in the study. Before inclusion, subjects underwent a medical examination where resting blood pressure, heart rate, and ECG of the subjects were measured. Furthermore, body composition was determined by dual-energy X-ray absorptiometry (Lunar DPX-IQ, GEHealthcare, Chalfont StGiles,UK). Inclusion criteria were 18–40 years of age and an active lifestyle, defined as more than 3 h of physical activity per week. Exclusion criteria were smoking, chronic disease, allergy towards medication, and the use of beta_2_-agonist or other prescription medication. Subjects were informed about risks and discomforts related to the study. Subject characteristics are presented in Table 1. Each subject gave written and oral informed consent prior to inclusion in the study. The study was approved by the Committee on Health Research Ethics of the Capital Region of Denmark (H-1-2012-119) and performed in accordance with the standards set by the Declaration of Helsinki. The study was registered in ClinicalTrials.gov (NCT02551276). Of the 13 subjects who were screened, 12 were included in the study of which all completed.

**Table 1.**
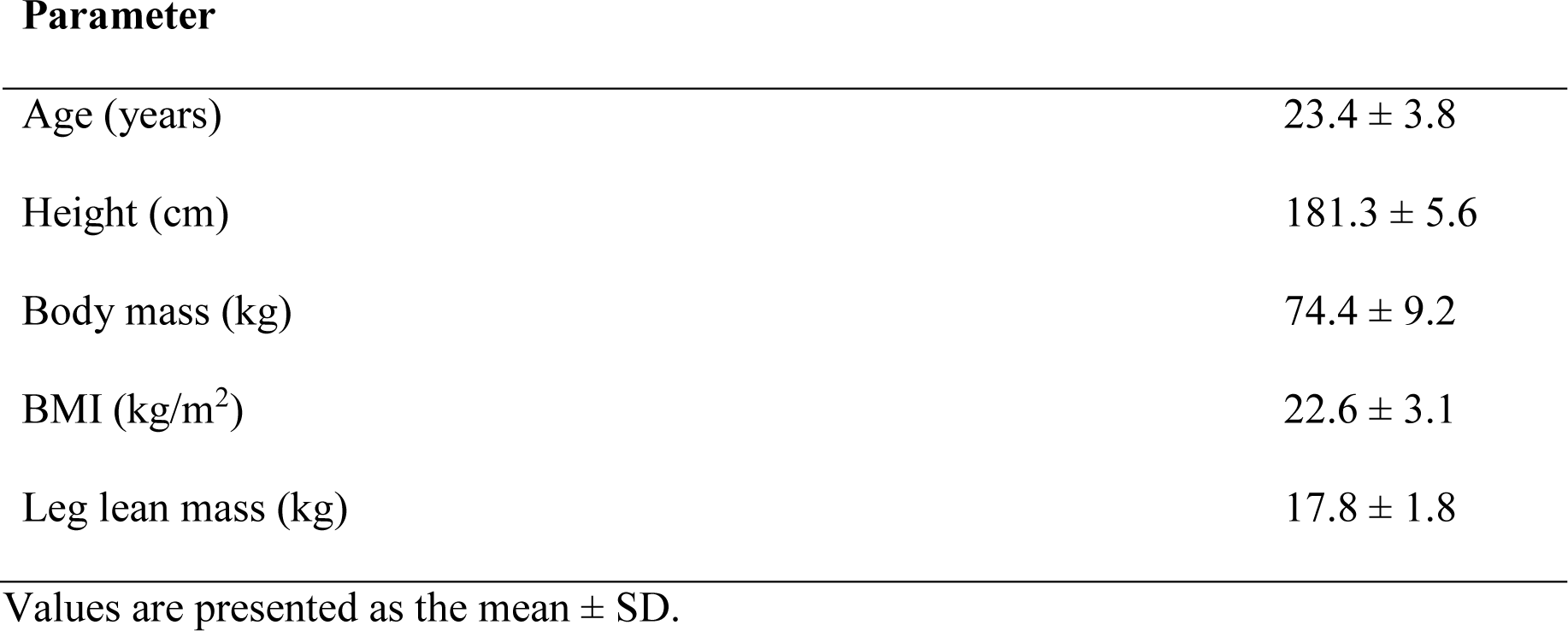
Subject characteristics (n = 12)

| Parameter |  |
| --- | --- |
| Age (years) | 23.4 ± 3.8 |
| Height (cm) | 181.3 ± 5.6 |
| Body mass (kg) | 74.4 ± 9.2 |
| BMI (kg/m<sup>2</sup>) | 22.6 ± 3.1 |
| Leg lean mass (kg) | 17.8 ± 1.8 |

Values are presented as the mean ± SD.

**Table 1.** Plasma insulin and muscle glycogen in recovery.

|  | Placebo |  | Salbutamol |  |
| --- | --- | --- | --- | --- |
|  | 0.5h | 5h | 0.5h | 5h |
| Insulin (pmol/L) | $62.8 \pm 7.7$ | $22.0 \pm 7.3$ | $76.3 \pm 7.7$ | $31.7 \pm 7.3^*$ |
| Glycogen (mmol/mg dry weight) | $295 \pm 18$ | $334^{\$} \pm 17$ | $186 \pm 18^*$ | $233^{\$} \pm 17^*$ |
| $\Delta$ glycogen (mmol/mg/dry weight) | | $39 \pm 13$ | | $47 \pm 15$ |

Values are presented as means ± SD. Plasma insulin concentration and muscle glycogen content at 0.5h and 5h into recovery for placebo and salbutamol. \*Treatment difference ( $P < 0.05$ ). \$Time difference ( $P < 0.05$ )

### Study design

This study was a part of a larger physiological study focusing on muscle metabolism in response to beta_2_-adrenergic stimulation and resistance exercise [20]. The study employed a randomized, double-blinded, placebo-controlled, cross-over design. During two identical experimental trials, subjects received either 24 mg salbutamol or placebo. Each trial was preceded by a 4-day lead-in period with oral salbutamol (4×4 mg/day) or placebo treatment because animal studies have shown that the effect of beta_2_-agonist on protein synthesis is evident after a few days of treatment [21]. The two trials were separated by 3–6 weeks to minimize potential confounding carry-over effects of salbutamol [22]. Before the first experimental trial, subjects met at the laboratory for two familiarizations to the resistance exercise protocol of the experimental trials.

### Experimental trials

After the lead-in treatment, subjects met in the morning after an overnight fast and received either oral salbutamol (6 × 4 mg) or placebo with a standardized light meal low on protein and fat consisting of white bread with jam (energy: 369 kcal; protein: 12 g; carbohydrate: 67 g; fat: 3 g) and 400 mL of water. Subjects then rested in a bed in the supine position and catheters were inserted: one in the dorsal hand vein, one in the brachial artery, and one in the femoral vein during local anesthesia (lidocaine without epinephrine, Xylocaine; AstraZeneca, Cambridge, UK) for arterial and venous blood sampling. A primed, continuous infusion of stable amino acid isotope [13C6]-phenylalanine (L-phenylalanine, ring-13C6, 99%, CLM-1055-MPT; Cambridge Isotope Laboratories, Inc., Tewksbury, MA, USA) was used for measurement of amino acid kinetics across the limb and incorporation of labeled phenylalanine into muscle as previously described [20]. The [13C6]-phenylalanine infusion equaled to an accumulated amount of ≈500-600 mg phenylalanine over 7 h, which is unlikely to affect substrate metabolism [23]. Furthermore, the infusion was conducted on both study days, why conditions for placebo and salbutamol trials were similar. Subjects then moved to a knee extensor resistance exercise model and performed two sets of 10 repetition knee extensor exercise at an intensity corresponding to 50% of three repetition maximum, followed by eight sets of 12 repetitions of knee-extensor exercise at an intensity corresponding to 12 repetition maximum (75 ± 11 kg) (mean ± SD) with 2 min of recovery between each set. If subjects failed to perform 12 repetitions in each set, the load was decreased for the following set. The mean load performed during the final set was 69 ± 12 kg. Intensity and recovery time were duplicated for each subject during the two trials. After exercise, subjects remained inactive in a supine position for 5 h. Biopsies were obtained from the vastus lateralis muscle 0.5 and 5 h after resistance exercise.

Brachial arterial and femoral venous blood samples were drawn in EDTA tubes (9 mL) before exercise as well as 0.5, 1, 2, 3, 4, and 5 h following exercise. Before exercise as well as 0.5, 1, 2, 3, 4, and 5 h following exercise, femoral arterial blood flow was measured with ultrasound Doppler (Vivid E9; GE Healthcare, Brøndbyvester, Denmark) equipped with a linear probe operating at an imaging frequency of 8 MHz and Doppler frequency of 3.1 MHz, as described previously [24]. Subjects were asked to refrain from caffeine, nicotine, and alcohol 24 h before each trial, as well as from exercise 48 h before each trial.

### Study drugs

Salbutamol (Ventolin, 4 mg tablets; GlaxoSmithKline, London, UK) and identically looking placebo (lactose monohydrate/starch) were delivered by the hospital pharmacy of Copenhagen. Drugs were administered in a double-blinded manner. Randomization was conducted in SPSS, version 24 (IBM Corp., Armonk, NY, USA) by personnel who did not take part in any of the experimental procedures or data analyses.

### Experimental procedures

#### Dual-energy X-ray absorptiometry

Subjects laid in the scanner in supine position undressed for 20 min before the scan. To reduce variation, two scans at medium speed were performed in accordance with the manufacturer’s guidelines. The scanner was calibrated before the scan using daily calibration procedures (Lunar ‘System Quality Assurance’). All scans were conducted by the same hospital technician.

#### Muscle biopsies

Muscle biopsies were obtained from the vastus lateralis using a 4-mm Bergström biopsy needle (Stille, Stockholm, Sweden) with suction (Bergström, 1975). Before biopsies were sampled, two incisions were made in the skin at the belly of the vastus lateralis muscle during local anesthesia (2 mL of lidocaine without epinephrine, Xylocaine 20 mg/mL; Astra Zeneca). After sampling, the muscle biopsy was cleaned from visible blood, connective tissue, and fat and immediately frozen in liquid nitrogen. Biopsies were stored in cryotubes at –80°C until analyses.

#### Immunoblotting

Protein content and degree of phosphorylation were determined by Western blotting as previously described [25]. Protein concentration of each sample was determined with a BSA kit (Thermo Fisher Scientific, Waltham, MA). Samples were mixed with 63 Laemmli buffer (7 mL 0.5M Tris-base, 3 mL glycerol, 0.93 g dithiothreitol, 1 g SDS, and 1.2 mg bromophenol blue) and double Q10 –distilled water to reach equal protein concentrations. Equal amounts of protein were loaded in each well of precast gels (Bio-Rad Laboratories, Hercules, CA). Samples from each subject were loaded on the same gel with a mixed human muscle standard lysate loaded in two different wells used for normalization. Bands were visualized with enhanced chemiluminescence (EMD Millipore, Billerica, MA) and recorded with a digital camera (Chemi- DocMP imaging system; Bio-Rad Laboratories). Densitometry quantification of band intensity was done using Image Lab version 4.0 (Bio-Rad Laboratories) and determined as the total band intensity adjusted for background intensity. Primary antibody used were for AMPKα2: a non-commercial antibody kindly donated by Graham Hardie, University of Dundee, 62 kDa; AMPKα^Thr172^: #2531 (Cell Signaling Technology), 62 kDa; PDH: sc - 377092 (Santa Cruz Biotechnology), 43 kDa; PDH^Ser293^: ABS204 (Millipore), 43 kDa; ACCβ: P-0397 (DAKO), 259 kDa; ACCβ^Ser221^: 07-303 (Millipore), 259 kDa, a phospho-specific ACCα^Ser79^ antibody recognizing the equivalent Ser^221^ in human ACCβ (Roepstorff *et al*. 2005 and Wojtaszewski *et al*. 2003). The secondary antibodies used were HRP conjugated goat anti-mouse, goat anti-rabbit, and rabbit anti-sheep (P-0447, P-0448 and P-0163 DAKO, Denmark).

#### Muscle glycogen content

Muscle glycogen was determined by the hexokinase method after extraction from dry frozen muscle tissue using 1NHCl for hydrolyzation at 100°C for 3 hours as previously described [26].

#### Blood sample analyses

Femoral arterial and venous blood samples were drawn in heparinized tubes for immediate analyses of glucose, lactate, hemoglobin, hematocrit, O_2_, CO_2_, and acid-base parameters using an ABL800 Flex (Radiometer, Copenhagen, Denmark). Further blood samples were drawn in EDTA tubes and stored on ice for 30 minutes before they were spun at 4°C and 3000 rpm for 5 minutes, after which plasma was collected and stored at −80°C until analysis for insulin. Plasma insulin was determined with ELISA (Dako Denmark) according to manufacturer’s instructions.

#### Calculations

Leg glucose and lactate exchange was calculated using Fick’s principle:

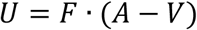

Where U is the net exchange of glucose or lactate, F is blood flow in the femoral artery, A is the arterial concentration and V is the femoral venous concentration.

#### Statistical analysis

Statistical analyses were performed in SPSS version 27 (IBM, Armonk, US). Data were tested for normality using the Shapiro-Wilks test and Q-Q plots. To estimate differences between treatments, a two-factor linear mixed modeling was used with treatment and time as fixed effects and a random effect for subjects. In addition, age and lean body mass were included in the model as time-invariant covariates because they may confound the effect of beta_2_-agonist [27,28]. Area under the uptake-/release-/clearance-time curve (AUC) was analyzed using the trapezoidal rule. Data are presented as mean with the 95% confidence interval (CI) and exact *p*-values (unless lower than 0.01) to represent probability for treatment fixed effects.

## Results

### Femoral arterial blood flow

With placebo, femoral arterial blood flow increased from resting values around 0.30 L/min to values of around 2.3 L/min during exercise and declined to values around 0.37 L/min in recovery, with the corresponding values being 181, 33, and 151% higher (P<0.001) for salbutamol, respectively (Fig. 1A & B).

**Figure 1.**
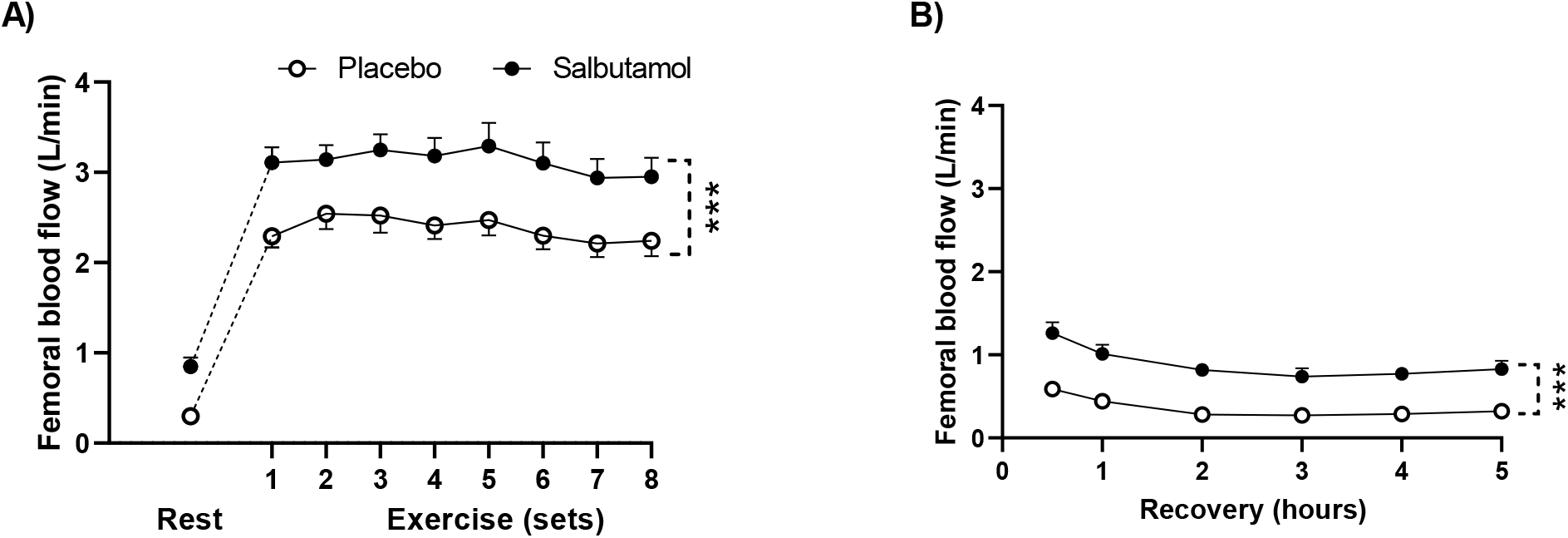
Femoral arterial blood flow at rest, during exercise and in recovery for placebo (white dots) and salbutamol (black dots). A) Femoral arterial blood flow at rest and during exercise. B) Femoral arterial blood flow in recovery. Values are presented as means ± SD. ***P < 0.001, for treatment difference.

### Leg glucose kinetics

While no difference was observed for arterial glucose concentration at rest (P=0.17), leg glucose delivery was approximately 5.0 ± 0.6 mmol/min higher (P<0.001) for salbutamol than placebo corresponding to an increase of 155%. Arterial glucose concentration and leg glucose delivery were 0.5 ± 0.1 mM and 5.2 ± 0.5mmol/min higher (P<0.001) for salbutamol than placebo during exercise equivalent to an increase of 10% and 45%, respectively. Likewise, arterial glucose concentration and leg glucose delivery were 0.5 ± 0.1 mM and 3.3 ± 0.4 mmol/min higher (P<0.001) for salbutamol than placebo in recovery amounting to an elevation of 10% and 170% (Fig. 2A & B). The femoral arteriovenous glucose difference was not different (P=0.27) between the treatments at rest, during exercise (P=0.63), or in recovery (P=0.41) (Fig. 2C & D). At rest, leg glucose uptake tended to be 0.22 ± 0.12 mmol/min for salbutamol (P=0.06) than placebo corresponding to a 112% increase, whereas leg glucose clearance was not different (P=0.13) with salbutamol compared to placebo. No treatment differences were observed for accumulated leg glucose uptake (P=0.49) or clearance (P=0.94) during exercise (Fig. 2E & 3A), but over the 5-h recovery, accumulated leg glucose uptake and clearance was 75 (P=0.018) and 65% higher (P=0.027)) for salbutamol than placebo (Fig. 2H & 3D).

**Figure 2:**
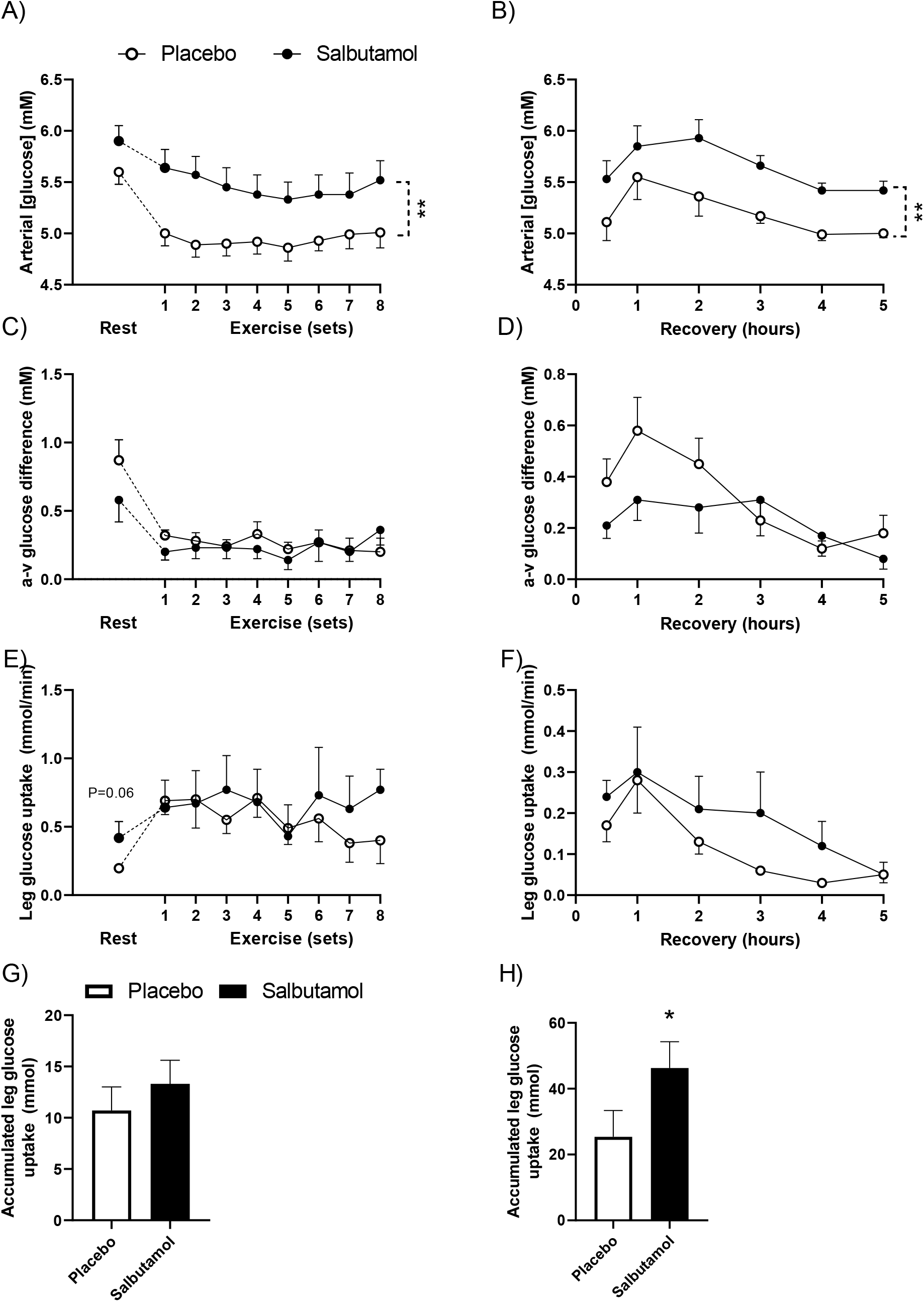
Leg glucose parameters at rest, during exercise and in recovery for placebo (white dots and bars) and salbutamol (black dots and bars). A) Femoral arterial glucose concentration at rest and during exercise. B) Femoral arterial glucose concentration in recovery. C) Femoral arteriovenous glucose difference at rest and during exercise. D) Femoral arteriovenous glucose difference in recovery. E) Leg glucose uptake at rest during exercise. F) Leg glucose uptake in recovery. G) Accumulated leg glucose uptake during exercise. H) Accumulated glucose uptake in recovery. Values are presented as means ± SD *P < 0.05, for treatment difference. **P < 0.001, for treatment difference.

**Figure 3:**
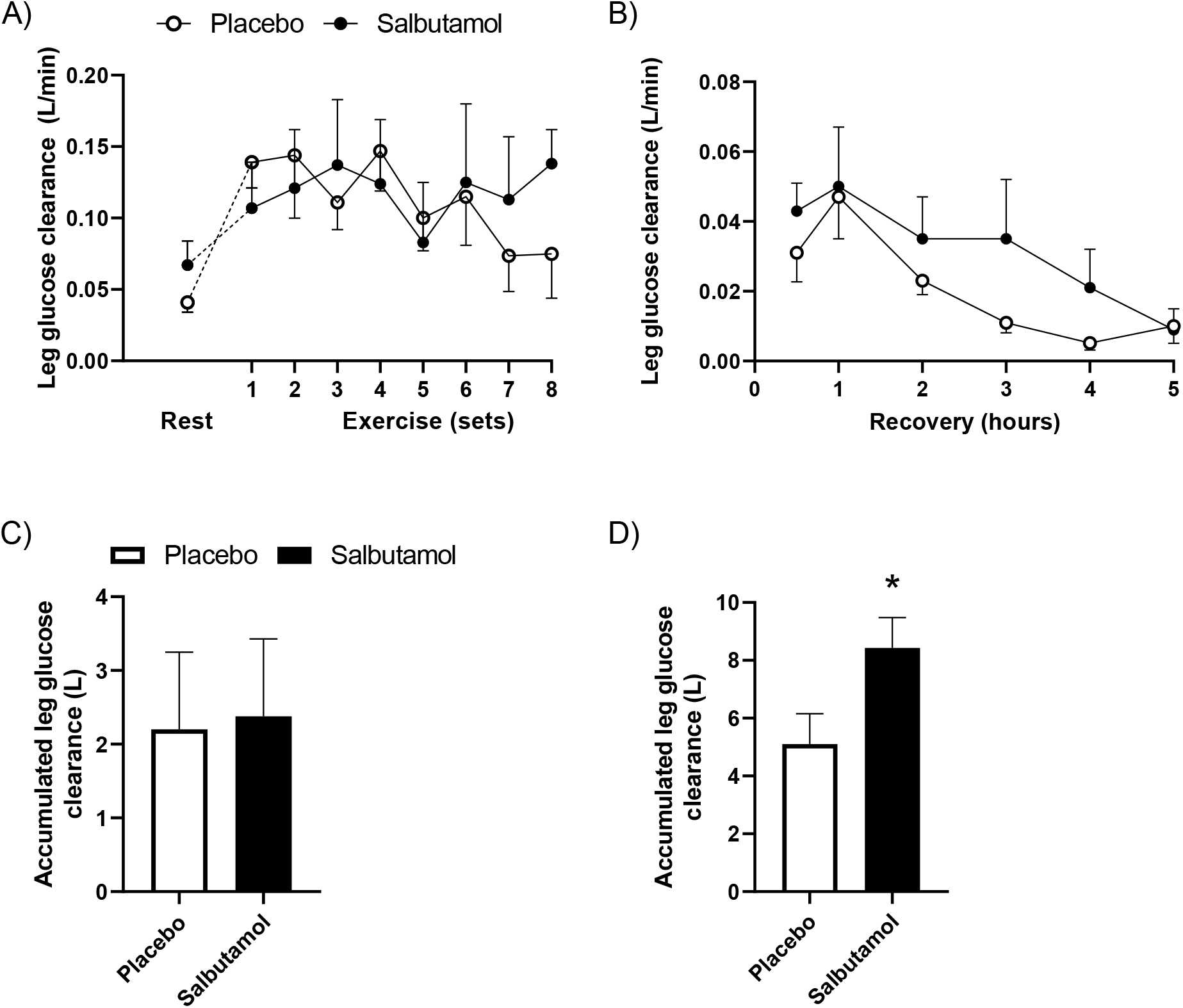
Leg glucose clearance and accumulated leg glucose clearance at rest, during exercise and in recovery for placebo (white dots and bars) and salbutamol (black dots and bars). A) Leg glucose clearance at rest and during exercise. B) Leg glucose clearance in recovery. C) Accumulated leg glucose clearance during exercise. D) Accumulated leg glucose clearance in recovery. Values are presented as means ± SD. *P < 0.05, for treatment difference.

### Leg lactate release

No treatment differences were observed in leg lactate release at rest (P=0.44)(Fig. 4A) nor in the accumulated amount of lactate released during exercise (P=0.49) (Fig. 4C), whereas net lactate release in recovery was 19.4 ± 10.3 mmol higher for salbutamol than placebo (P < 0.01) (Fig. 4D) corresponding to a 4-fold increase.

**Figure 4:**
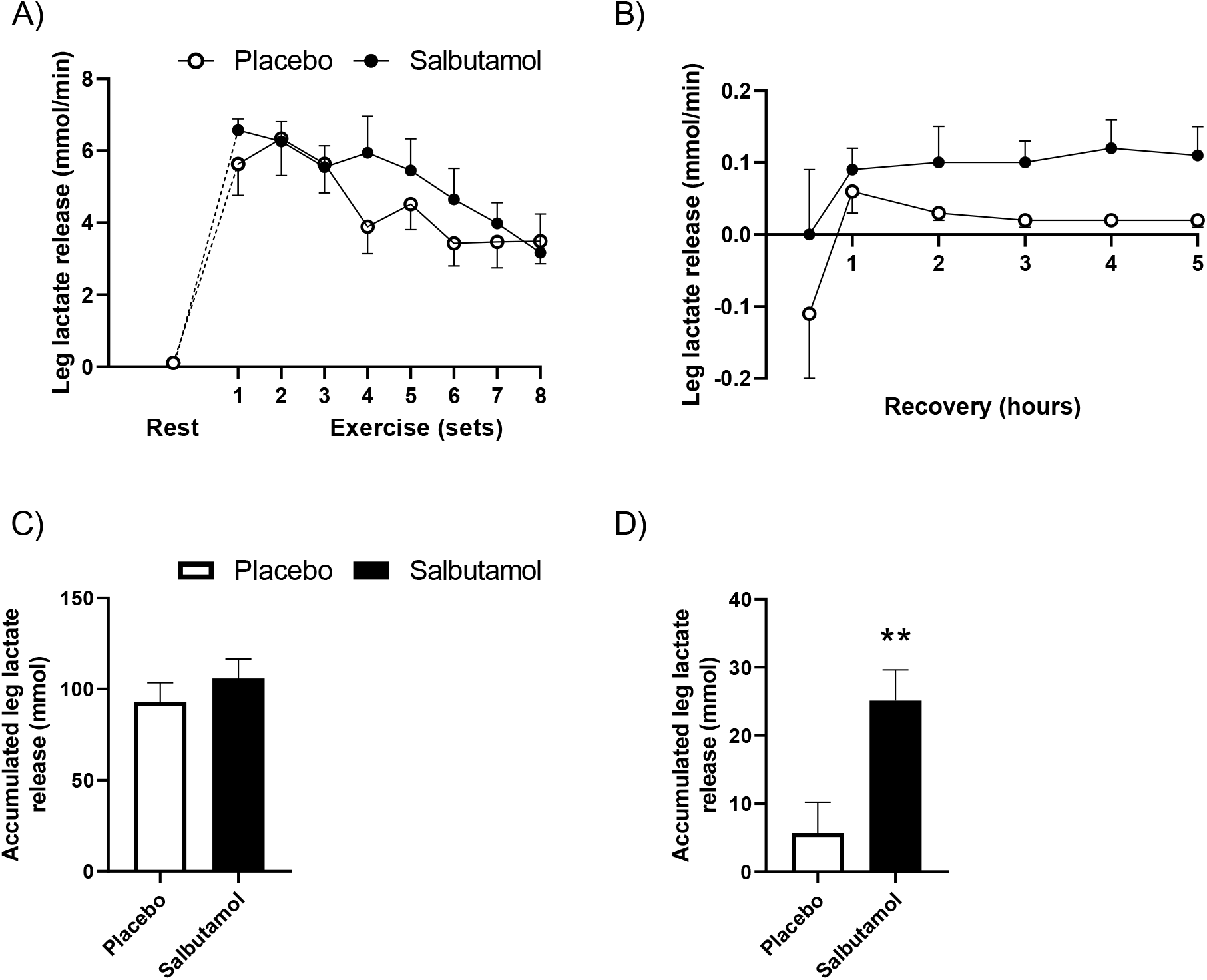
Leg lactate release and accumulated leg lactate release at rest, during exercise and in recovery for placebo (white dots and bars) and salbutamol (black dots and bars). A) Leg lactate release at rest and during exercise. B) Leg lactate release in recovery. C) Accumulated leg lactate release during exercise. D) Accumulated leg lactate release in recovery. Values are presented as means ± SD. **P < 0.01, for treatment difference.

### Leg oxygen uptake

Leg oxygen uptake was 80% greater with salbutamol than placebo at rest (P<0.005), The accumulated amount of oxygen extracted by the leg was not different between treatments during exercise (P=0.24)(Fig. 5B) but was 1755 ± 825 mL higher in recovery for salbutamol than placebo (P < 0.01)(Fig. 5D).

**Figure 5:**
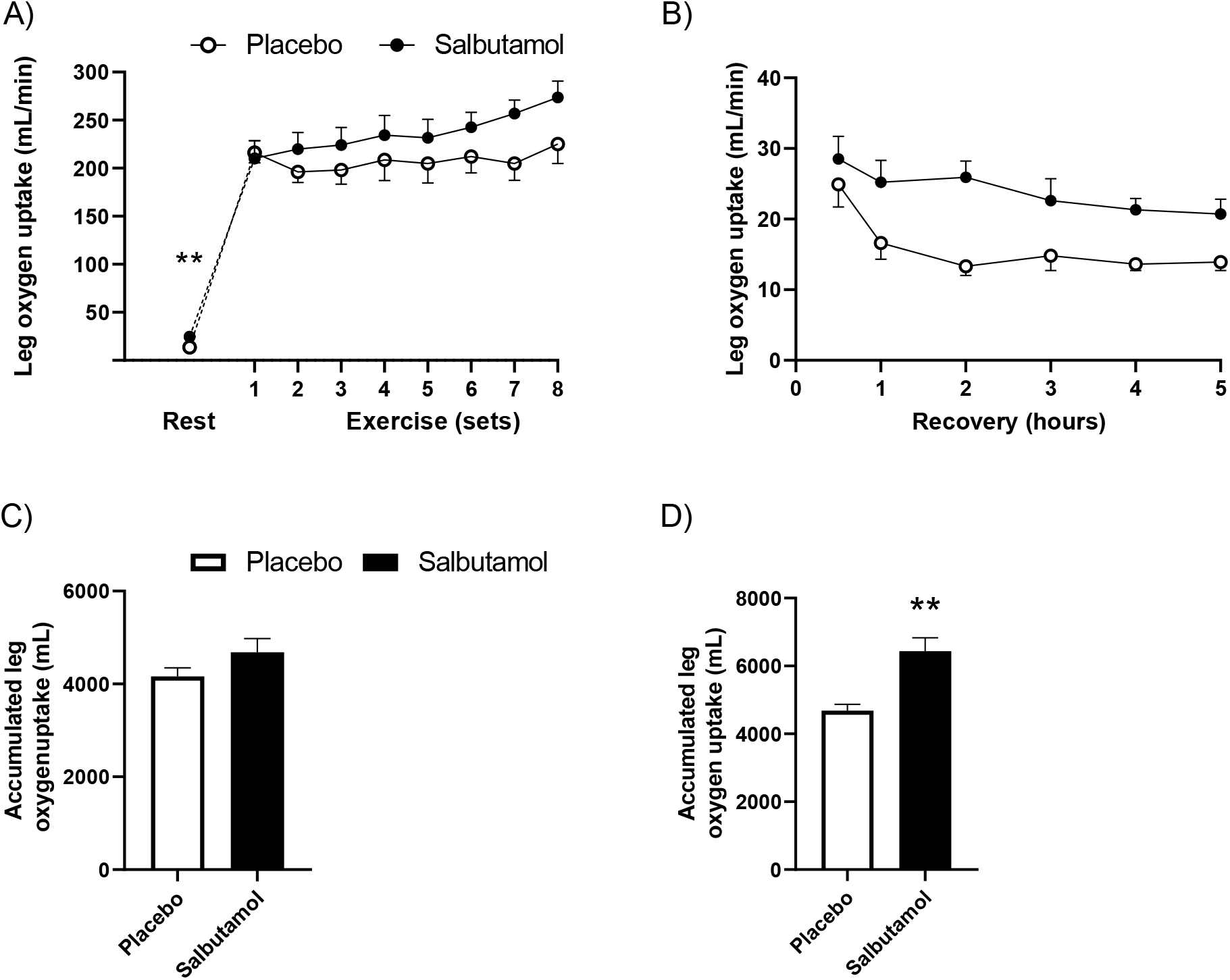
Leg oxygen uptake and total leg oxygen uptake at rest, during exercise and in recovery for placebo (white dots and bars) and salbutamol (black dots and bars). A) Leg oxygen uptake during exercise. B) Leg oxygen uptake in recovery. C) Accumulated leg oxygen uptake during exercise. D) Accumulated leg oxygen uptake in recovery. Values are presented as means ± SD. ** P < 0.01, for treatment difference. ** P< 0.001 for treatment difference.

### Plasma insulin concentration

The arterial plasma insulin concentration was not significantly different (P=0.11) between the treatments 0.5 h into recovery (Table 1) but was 44% higher (P=0.03) for salbutamol than placebo 5 h into recovery (Table 1).

### Muscle glycogen content and muscle glycogen synthase activity

Muscle glycogen content was 37% and 51% lower for salbutamol than placebo 0.5 and 5 h into recovery, respectively (P < 0.01) (Table 1). Over the recovery period from 0.5 to 5 h, muscle glycogen content increased by 24% for salbutamol (P<0.01), which was not different from the increase of 13% for placebo (P<0.01) (Table 1). There was no difference in muscle glycogen synthase activity between placebo and salbutamol at either 0.5 h or 5 h into the recovery (P>0.5) (Fig. 6).

**Figure 6:**
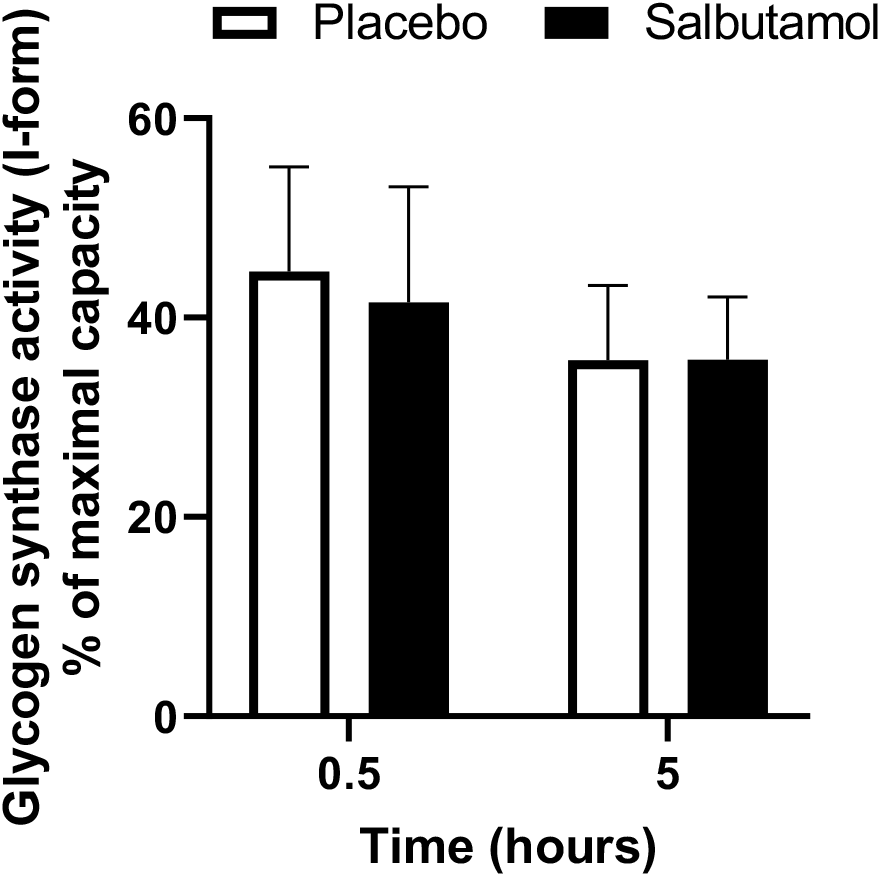
Glycogen synthase activity measured as fractional activity in the presence of 0.2 mM G6P and given relative to saturated conditions (8 mM G6P) at time points 0.5 h and 5 h into recovery for placebo (white bars) and salbutamol (black bars). Values are presented as means ± SD. ^$^P < 0.05 for time difference.

### Muscle AMPK, PDH, and ACCβ phosphorylation

Muscle AMPK^Thr172^ phosphorylation was 22% higher (P=0.038) for salbutamol than placebo 0.5 h but not 5 h into recovery (P=0.94) (Fig. 7A). No treatment difference was observed in AMPKα2 at either 0.5 h or 0.5 h into recovery (P=0.60 and P=0.32, respectively) (Fig. 7B). Given there is no available anti-body to quantify total AMPK, AMPKα2 has been measured instead. As both AMPKα2 and AMPKα1 are phosphorylated on threonine 172, a ratio between total AMPK and AMPKα2 has not been calculated. Muscle PDH^Ser293^ phosphorylation was 30 and 20% higher 0.5 and 5 h into recovery (p < 0.01) for salbutamol than placebo, respectively (Fig. 7C), while phosphorylation ratio PDH^Ser293^ / total PDH was 20 and 15% higher 0.5 and 5 h into recovery (P < 0.01) for salbutamol than placebo, respectively (Fig. 7D). Muscle ACCβ^Ser221^ phosphorylation was 75% higher (P=0.048) 0.5h into recovery (Fig. 7E) and 2.4 fold higher (P < 0.01) 5 h into recovery (Fig. 7E) for salbutamol than placebo. Phosphorylation ratio ACCβ^Ser221^ / total ACCβ was not different between treatments 0.5 h in recovery (P=0.33) (Fig. 7F), but was 83% higher (P < 0.01) for salbutamol than placebo 5 h into recovery (Fig. 7F). Representative blots are shown in figure 7.

**Figure 7:**
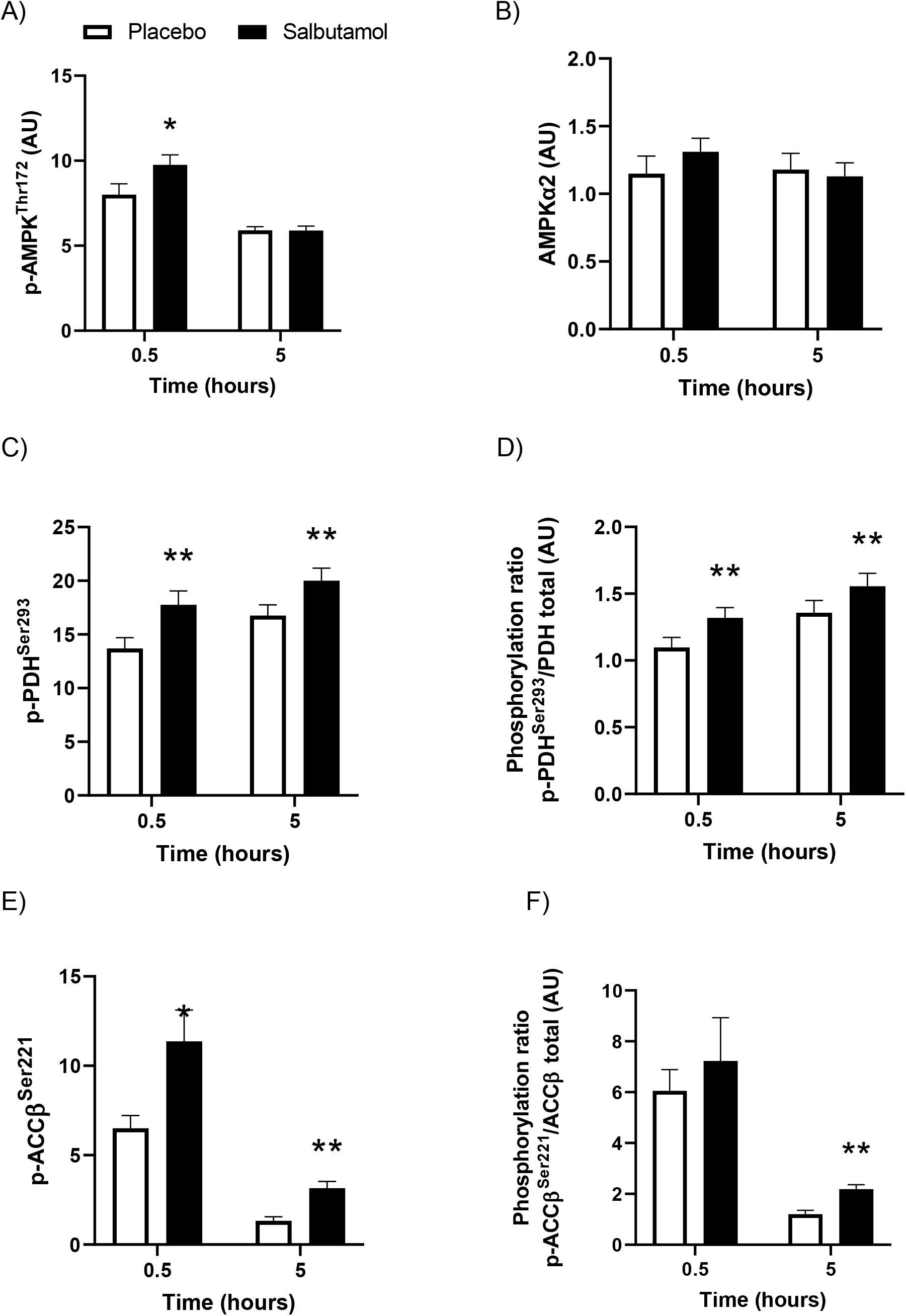
Muscle protein phosphorylation, phosphorylation ratios and AMPKα2 at time points 0.5 h and 5 h into recovery for placebo (white bars) and salbutamol (black bars). Represented as both phosphorylation (A, C, E) and phosphorylation compared to total protein (B, D, F). A) Phosphorylation of AMPK (Threonine-172). B) AMPKα2. C) Phosphorylation of PDH (Serine-293). D) Phosphorylation ratio of PDH (Serine-293) / Total PDH. E) Phosphorylation of ACC β(Serine 221). F) Phosphorylation ratio of ACC β(Serine 221) / total ACC_β_. Values are presented as means ± SD. *P < 0.05, ** P < 0.01 for treatment difference.

## Discussion

The major finding of this study was that beta_2_-agonist salbutamol augmented leg glucose uptake at rest and in recovery from work-matched resistance exercise of the quadriceps in lean young men. This effect, however, was blunted during the exercise. Despite the greater glucose uptake in recovery, salbutamol did not augment muscle glycogen resynthesis compared to placebo. In fact, muscle glycogen levels were lower for salbutamol 0.5-5 h after the exercise than for placebo. Additionally, the rate of lactate release and amount of oxygen extracted from the exercised leg was higher for salbutamol, underpinning the thermogenic action of beta_2_-agonists in skeletal muscle. These salbutamol-induced changes were accompanied by greater muscle phosphorylation of PDH, AMPK, and ACCβ in recovery.

The observation that salbutamol led to a marked increase of leg glucose uptake and clearance in recovery from work-matched exercise, but not during exercise, extends previous studies by showing that the confounding effect of exercise on the beta_2_-agonist-induced augmentation of leg glucose uptake diminishes during recovery. Hence, the 75% greater leg glucose uptake in recovery with salbutamol is strikingly similar to the two-fold increase in leg glucose uptake observed at pre-exercise resting conditions in the present study and in studies utilizing terbutaline infusion [9]. While one might speculate that this effect could be due to a greater glucose delivery with salbutamol, this did not seem to be the case as leg glucose clearance was still 65% greater for salbutamol than placebo in recovery. Thus, the bulk of the augmented glucose disposal with beta_2_-agonist appears dissociated from its effects on glucose availability and may instead be related to the higher metabolic rate of the exercised leg in recovery with salbutamol, as indicated by the greater leg oxygen extraction and lactate release for salbutamol than placebo.

Other factors underpinning the greater glucose uptake could relate to the effect of salbutamol on insulin levels and canonical adrenergic muscle PKA-signalling. However, given that leg glucose uptake was four times greater for salbutamol than placebo 4 h into recovery, where insulin was only 44% higher for salbutamol, the hyperinsulinemic effect of salbutamol possibly only partly contributed to the augmented leg glucose uptake. Indeed, Sato et al. observed that beta_2_-adrenergic stimulation of myotubes potently increased glucose uptake through canonical adrenergic PKA-signaling, and independent of insulin-signalling [10]. The augmented leg glucose uptake in recovery could also be related to the markedly lower muscle glycogen levels after the work-matched exercise for salbutamol than placebo. Indeed, activation of AMPK is associated with the level of glycogen depletion [17] and AMPK contributes to the regulation of myocellular glucose uptake in recovery from exercise [18]. Hence, salbutamol could facilitate a more prominent AMPK-activation, which is supported by the 22% greater phosphorylation of muscle AMPK ^thr172^ for salbutamol than placebo ½ h into recovery observed in the present study. Further indicative of greater AMPK activation, ACCβ, a downstream target of AMPK, was 75% more phosphorylated ½ h into recovery for salbutamol than placebo and more than two-fold more phosphorylated 5 h into recovery.

A notable 4-fold higher amount of lactate was released from the exercised leg in recovery for salbutamol than placebo. Because of the absence of treatment differences in glycogen resynthesis, this is indicative of pyruvate accumulating in the glycolysis to be shunted towards lactate formation with salbutamol. Indeed, phosphorylation of PDH^ser293^ was potently amplified by 30% for salbutamol compared to placebo in recovery, suggesting a deactivation of PDH. Hence, one could imagine a scenario in which excessive amounts of glucose are taken up in recovery with salbutamol, which due to deactivation of PDH, is directed towards lactate production. Furthermore, despite the markedly lower glycogen levels in recovery from exercise with salbutamol, this did not facilitate a greater activity of glycogen synthase nor in the amount of glycogen resynthesized during the 4½ h period for salbutamol than placebo. This suggests that the expected increase in the activity of glycogen synthase facilitated by a greater glycogen depletion with salbutamol was negated, likely via an inhibitory effect of beta_2_-adrenergic stimulation on glycogen synthase as observed in other studies [19]. Such inhibition would also contribute to the greater rate of lactate production observed with salbutamol in recovery.

Consistent with previous studies [9], we observed no apparent effect of salbutamol on leg glucose uptake during exercise. The observation that exercise can blunt the effect of beta_2_-agonist is possibly related to the cascade of intrinsic events induced by exercise in skeletal muscle to augment glucose uptake via translocation and redistribution of GLUT4 as being sufficiently large to mask the effect of beta_2_-agonist. Furthermore, the bulk of the effect of beta_2_-agonist on leg glucose uptake at resting conditions seems to be due to the greater muscle metabolic rate and to a lesser extent excess glucose delivery. While the latter was still apparent during exercise, salbutamol had no apparent effect on leg metabolic rate during the work-matched exercise, in terms of leg VO_2_ and net lactate release. This again supports that it is the effect of beta_2_-agonist on muscle metabolic rate that explains the majority of the effect observed on leg glucose uptake rather than glucose delivery *per se*.

Taken together, these findings confirm that exercise can confound the effect of beta_2_-agonist on leg glucose uptake but that this attenuation is diminished in recovery. Our findings also suggest that the greater leg lactate release induced by salbutamol in recovery from exercise may be due to glycolytic trafficking due to excessive myocellular glucose uptake concomitant with deactivation of PDH. Considering leg glucose uptake and metabolic rate are preserved in recovery from exercise, the current study underlines the potential of the beta_2_-adrenoceptor as a future therapeutic target in treatment of T2DM and obesity. Furthermore, the pronounced effects on muscle metabolism of the oral doses of salbutamol administered support the restrictions imposed on salbutamol in sports.

## References

[1] Price OJ, Hull JH, Backer V, Hostrup M, Ansley L. The Impact of Exercise-Induced Bronchoconstriction on Athletic Performance: A Systematic Review. Sport Med 2014;44:1749–61. https://doi.org/10.1007/s40279-014-0238-y.

[2] Kainu A, Pallasaho P, Piirilä P, Lindqvist A, Sovijärvi A, Pietinalho A. Increase in prevalence of physician-diagnosed asthma in Helsinki during the Finnish Asthma Programme: improved recognition of asthma in primary care? A cross-sectional cohort study. Prim Care Respir J 2013;22:64–71. https://doi.org/10.4104/pcrj.2013.00002.

[3] Lynch GS, Ryall JG. Role of b-Adrenoceptor Signaling in Skeletal Muscle: Implications for Muscle Wasting and Disease. Physiol Rev 2008;88:729–67. https://doi.org/10.1152/physrev.00028.2007.

[4] Jensen J, Brennesvik EO, Bergersen L, Oseland H, Jebens E, Brørs O. Quantitative determination of cell surface β-adrenoceptors in different rat skeletal muscles. Pflügers Arch 2002;444:213–9. https://doi.org/10.1007/s00424-002-0793-1.

[5] Schiffelers SLH, Saris WHM, Boomsma F, van Baak MA. β _1_- andβ_2_-Adrenoceptor-Mediated Thermogenesis and Lipid Utilization in Obese and Lean Men ^1^. J Clin Endocrinol Metab 2001;86:2191–9. https://doi.org/10.1210/jcem.86.5.7506.

[6] Hostrup M, Kalsen A, Ørtenblad N, Juel C, Mørch K, Rzeppa S, et al. β2-Adrenergic stimulation enhances Ca2+ release and contractile properties of skeletal muscles, and counteracts exercise-induced reductions in Na+-K+-ATPase Vmax in trained men. J Physiol 2014;592:5445–59. https://doi.org/10.1113/jphysiol.2014.277095.

[7] van Baak MA, de Haan A, Saris WH, van Kordelaar E, Kuipers H, van der Vusse GJ. Beta-adrenoceptor blockade and skeletal muscle energy metabolism during endurance exercise. J Appl Physiol 1995;78:307–13. https://doi.org/10.1152/jappl.1995.78.1.307.

[8] Onslev J, Jacobson G, Narkowicz C, Backer V, Kalsen A, Kreiberg M, et al. Beta2-adrenergic stimulation increases energy expenditure at rest, but not during submaximal exercise in active overweight men. Eur J Appl Physiol 2017;117:1907–15. https://doi.org/10.1007/s00421-017-3679-9.

[9] Onslev J, Jensen J, Bangsbo J, Wojtaszewski J, Hostrup M. β 2-Agonist Induces Net Leg Glucose Uptake and Free Fatty Acid Release at Rest but Not during Exercise in Young Men. J Clin Endocrinol Metab 2018;104:647–57. https://doi.org/10.1210/jc.2018-01349.

[10] Sato M, Dehvari N, Öberg AI, Dallner OS, Sandström AL, Olsen JM, et al. Improving type 2 diabetes through a distinct adrenergic signaling pathway involving mTORC2 that mediates glucose uptake in skeletal muscle. Diabetes 2014;63:4115–29. https://doi.org/10.2337/db13-1860.

[11] Kalinovich A, Dehvari N, Åslund A, van Beek S, Halleskog C, Olsen J, et al. Treatment with a β-2-adrenoceptor agonist stimulates glucose uptake in skeletal muscle and improves glucose homeostasis, insulin resistance and hepatic steatosis in mice with die t-induced obesity. Diabetologia 2020;63:1603–15. https://doi.org/10.1007/s00125-020-05171-y.

[12] Lund J, Gillum MP. Towards Leanness by ‘Feeding’ a Novel Thermogenic Pathway? Trends Endocrinol Metab 2016;27:529–30. https://doi.org/10.1016/j.tem.2016.05.006.

[13] Arlettaz A, Le Panse B, Portier H, Lecoq A -M, Thomasson R, De Ceaurriz J, et al. Salbutamol intake and substrate oxidation during submaximal exercise. Eur J Appl Physiol 2009;105:207–13.

[14] Kalsen A, Hostrup M, Karlsson S, Hemmersbach P, Bangsbo J, Backer V. Effect of inhaled terbutaline on substrate utilization and 300-kcal time trial performance. J Appl Physiol 2014;117:1180–7. https://doi.org/10.1152/japplphysiol.00635.2014.

[15] Praet SFE, Van Loon LJC. Exercise therapy in Type 2 diabetes. Act a Diabetol 2009;46:263–78. https://doi.org/10.1007/s00592-009-0129-0.

[16] Background A. Expert panel report: Guidelines (2013) for the management of overweight and obesity in adults. Obesity 2014;22:S41–410. https://doi.org/10.1002/oby.20660.

[17] Wojtaszewski JFP, MacDonald C, Nielsen JN, Hellsten Y, Grahame Hardie D, Kemp BE, et al. Regulation of 5′-AMP-activated protein kinase activity and substrate utilization in exercising human skeletal muscle. Am J Physiol - Endocrinol Metab 2003;284. https://doi.org/10.1152/ajpendo.00436.2002.

[18] Kjøbsted R, Roll JLW, Jørgensen NO, Birk JB, Foretz M, Viollet B, et al. AMPK and TBC1D1 regulate muscle glucose uptake after, but not during, exercise and contraction. Diabetes 2019;68:1427–40. https://doi.org/10.2337/db19-0050.

[19] Jensen J, Ruge T, Lai YC, Svensson MK, Eriksson JW. Effects of adrenaline on whole-body glucose metabolism and insulin-mediated regulation of glycogen synthase and PKB phosphorylation in human skeletal muscle. Metabolism 2011;60:215–26. https://doi.org/10.1016/j.metabol.2009.12.028.

[20] Hostrup M, Reitelseder S, Jessen S, Kalsen A, Nyberg M, Egelund J, et al. Beta2-adrenoceptor agonist salbutamol increases protein turnover rates and alters signalling in skeletal muscle after resistance exercise in young men. J Physiol 2018;596:4121–39. https://doi.org/10.1113/JP275560.

[21] Koopman R, Gehrig SM, Léger B, Trieu J, Walrand S, Murphy KT, et al. Cellular mechanisms underlying temporal changes in skeletal muscle protein synthesis and breakdown during chronic β-adrenoceptor stimulation in mice. J Physiol 2010;588:4811–23. https://doi.org/10.1113/jphysiol.2010.196725.

[22] Le Panse B, Collomp K, Portier H, Lecoq AM, Jaffre C, Beaupied H, et al. Effects of short - term salbutamol ingestion during a Wingate test. Int J Sports Med 2005;26:518–23. https://doi.org/10.1055/s-2004-821224.

[23] Holm L, Reitelseder S, Dideriksen K, Nielsen RH, Bülow J, Kjaer M. The single-biopsy approach in determining protein synthesis in human slow-turning-over tissue: Use of flood-primed, continuous infusion of amino acid tracers. Am J Physiol - Endocrinol Metab 2014;306:1330–9. https://doi.org/10.1152/ajpendo.00084.2014.

[24] Nyberg M, Christensen PM, Mortensen SP, Hellsten Y, Bangsbo J. Infusion of ATP increases leg oxygen delivery but not oxygen uptake in the initial phase of intense knee - extensor exercise in humans. Exp Physiol 2014;99:1399–408. https://doi.org/10.1113/expphysiol.2014.081141.

[25] Thomassen M, Christensen PM, Gunnarsson TP, Nybo L, Bangsbo J. Effect of 2-wk intensified training and inactivity on muscle Na+-K+ pump expression, phospholemman (FXYD1) phosphorylation, and performance in soccer players. J Appl Physiol 2010;108:898–905. https://doi.org/10.1152/japplphysiol.01015.2009.

[26] Lowry O, Passonneau J. A flexible system of enzymatic analysis. New York: Academic Press; n.d.

[27] Cheymol G. Effects of obesity on pharmacokinetics implications for drug therapy. Clin Pharmacokinet 2000;39:215–31. https://doi.org/10.2165/00003088-200039030-00004.

[28] White M, Leenen FHH. Aging and cardiovascular responsiveness to β-agonist in humans: Role of changes in β-receptor responses versus baroreflex activity. Clin Pharmacol Ther 1994;56:543–53. https://doi.org/10.1038/clpt.1994.176.

